# Association of Bile Acids with Connectivity of Executive Control and Default Mode Networks in Patients with Major Depression

**DOI:** 10.1101/2024.12.20.629637

**Authors:** Boadie W. Dunlop, Jungho Cha, Helen S. Mayberg, Ki Sueng Choi, W. Edward Craighead, Siamak MahmoudianDehkordi, Sudeepa Bhattacharyya, A. John Rush, Rima Kaddurah-Daouk

## Abstract

**Objective:** Bile acids may contribute to pathophysiologic markers of Alzheimer’s disease, including disruptions of the executive control network (ECN) and the default mode network (DMN). Cognitive dysfunction is common in major depressive disorder (MDD), but whether bile acids impact these networks in MDD patients is unknown.

**Methods:** Resting state functional magnetic resonance imaging (fMRI) scans and blood measures of four bile acids from 74 treatment-naïve adults with MDD were analyzed. Dorsolateral prefrontal cortex (DLPFC) seeds were used to examine connectivity of the ECN and posterior cingulate cortex (PCC) seeds were used for the DMN. Using a whole-brain analysis, the functional connectivity of these seeds was correlated with serum levels chenodeoxycholic acid (CDCA) and its bacterially-derived secondary bile acid, lithocholic acid (LCA).

**Results:** CDCA levels were strongly and inversely correlated with connectivity between DLPFC regions of the ECN (R^2^= .401, p<.001). LCA levels were strongly and positively correlated with connectivity of the DLPFC and left inferior temporal cortex of the ECN (R^2^=.263, p<.001). The LCA/CDCA ratio was strongly and positively correlated with connectivity of the DLPFC with two components of the ECN: bilateral inferior temporal cortex and the left superior and inferior parietal lobules (all R^2^>.24, all p<.001). For the DMN, the LCA/CDCA ratio was strongly and negatively correlated with connectivity of the PCC with multiple bilateral insula regions (all R^2^>0.25, all p<.001).

**Conclusions:** The relationship between LCA and CDCA levels and functional connectivity of the ECN and DMN suggests potential shared pathophysiologic processes between Alzheimer’s disease and MDD.

## INTRODUCTION

Major depressive disorder (MDD) is a common psychiatric disorder with substantial heterogeneity in symptom presentation. Although this heterogeneity has been the focus of much study, understanding of the biological underpinnings for this variation remain very limited. It is likely that the syndrome of MDD stems from a variety of pathobiological disturbances. This presumed multiplicity in pathophysiological mechanisms contributes to the high variability of response to treatments for MDD, including psychotherapy, medication, and stimulation treatments (1). Greater understanding of the biological alterations across MDD patients could help achieve the goals of “precision medicine” so that treatments administered will address specific biological targets (2).

Impairments in cognitive function are common but not universal in patients with MDD (3). One quarter of MDD patients demonstrate objective cognitive impairment while nearly two-thirds have reductions in cognitive function compared to their estimated premorbid level of function (4). The cognitive domains most affected in MDD include executive functioning, concentration, working memory, and sustained attention (5). These impairments, though more common in older depressed adults, also occur in younger patients with MDD (6). The presence of cognitive impairments predicts poorer treatment outcomes (7) and greater disability in both occupational (8) and social (9) settings. Impaired cognitive functioning may continue even after achieving symptomatic remission and it tends to worsen with repeated major depressive episodes (10). Experiencing a major depressive episode in one’s lifetime more than doubles the risk of developing dementia later in life, even when the episode occurred more than 20 years prior to a dementia diagnosis (11). Potential mechanisms underlying this association include shared genetic risks, social isolation stemming from MDD, or inflammatory processes.

Insights from mapping intrinsic brain connectivity networks provide a mechanistic framework for an understanding of aspects of human behavior (12). Cognitive functioning is well-established to be dependent upon activity within the brain’s executive control network (ECN, also known as the control network or frontal-parietal network). Key components of this network include the dorsolateral prefrontal cortex (DLPFC), posterior parietal cortex, and posterior-inferior temporal cortex. Impaired or inefficient functioning of the ECN in MDD underlies impairments in executive function, working memory, and attention that are needed for addressing demanding cognitive tasks (13). Multiple studies of healthy adults and MDD patients have demonstrated correlations between impairments in connectivity within this network and poorer performance on cognitive tasks (14,15). Resting state functional magnetic imaging (rs-fMRI) studies of AD and mild cognitive impairment (MCI) patients have also implicated abnormalities in the ECN, along with other brain networks, including the default mode network (DMN) (16). The DMN’s primary components are the posterior cingulate/medial parietal cortex and the medial prefrontal cortex. DMN activity in MDD is elevated compared to healthy controls (17) and is thought to be specifically involved in ruminative thought processes (18).

Recent studies in both MDD and MCI/AD have implicated alterations in bile acids (BA) as potential causal factors for neuropsychiatric diseases, potentially related to their effects on cholesterol metabolism or inflammation (19). BA were long considered to function purely as surfactants to support absorption of fats and lipid-soluble substances from the intestine. It is now clear that BA have functions across a variety of physiologic processes, including those relevant to brain function. After being produced in the liver, BA undergo differential metabolism, depending largely upon the composition of the individual’s gut microbiome. Reviews of gut microbiome studies in MDD have found inconsistent results (20,21), leading to the recommendation that the study of microbial functioning and metabolite production may be moree productive than a focus on bacterial species and genera, given that bacterial functions are conserved across taxonomic groups (20). BAs produced via the gut microbiome can impact the experience of anxiety and depression by various mechanisms, including activation of afferent vagal nerve fibers, stimulation of the mucosal immune system or circulatory immune cells after translocation from the gut into the circulation, and absorption into the bloodstream followed by uptake and biochemical interaction with a number of distal organs (22). In the brain, these metabolites may activate receptors on neurons or glia and modulate neuronal excitability, thereby changing gene expression patterns via epigenetic mechanisms (23).

Hepatocytes convert cholesterol to cholic acid (CA) and chenodeoxycholic acid (CDCA), the two most common primary BAs. After secretion into the small intestine, most (roughly 95%) of the CA and CDCA are resorbed in the ileum (19), but a small proportion enters the colon where bacterially-mediated 7α-dehydroxylation converts CA to deoxycholic acid (DCA) and CDCA to lithocholic acid (LCA); DCA and LCA are the most common secondary BAs (24).

Although production of primary BAs by glia and astrocytes can occur, the great majority of primary BAs and all secondary BAs in the brain are believed to enter via diffusion or active transport from the systemic circulation (22). BA concentrations in serum positively correlate with brain concentrations (25), allowing use of sampling of blood BAs to serve as a reasonable proxy for central nervous system BA concentrations.

Multiple studies have identified a role for BAs in cognitive dysfunction in CNS diseases, including hepatic encephalopathy (26) and AD (27–29). Patients with MDD demonstrate plasma concentrations of conjugated forms of LCA that significantly differ from those in health controls (30). Although secondary BA profiles reflective of gut microbiome compositions have been associated with higher levels of anxiety and poorer response to depression treatments (31), the impact of BAs on the neural functioning in psychiatric diseases such as MDD has received little study to date. We theorized that BA-associated pathology in AD may be present in patients with MDD and correlate with intrinsic brain network functioning.Here, using a cross-sectional design, we evaluated associations between pre-treatment baseline serum BA concentrations and rs-fMRI functional connectivity within the ECN and DMN among patients enrolled in a randomized controlled trial of treatment-naïve outpatients with MDD.

## METHODS

### Study Design and Participants

The Predictors of Remission in Depression to Individual and Combined Treatments (PReDICT) study was designed to identify predictors and moderators of response to 12 weeks of randomly assigned treatment with antidepressant medication or cognitive behavior therapy (CBT). Details about the design (32), clinical outcomes (33) and neuroimaging results (34–36) of PReDICT are published elsewhere. The current analysis focused on the baseline (pre-treatment) associations between peripheral bile acid levels and neuroimaging data in the participants. Eligible participants were adult outpatients aged 18-65 who met criteria based on the Structured Clinical Interview for DSM-IV (37) for MDD, single episode or recurrent, without psychotic features, who reported no lifetime treatment with medication or an evidence-based psychotherapy for depression. Exclusion criteria included a personal lifetime history of bipolar disorder, anorexia nervosa, a neurocognitive disorder or history of stroke, or current significant suicide risk, pregnancy, or lactation, or had current illicit drug use or a history of substance abuse in the three months prior to baseline, or any uncontrolled general medical condition. All patients provided written informed consent to participate. The study was approved by Emory University’s Institutional Review Board and the Grady Hospital Research Oversight Committee.

### Clinical Assessments

Severity of depression was assessed with the 17-item Hamilton Depression Rating Scale (HRSD_17_) (38), which has a range or 0-52. A score ≥15 (indiciating moderate depression) required for randomization at the baseline visit. Anxiety at baseline was assessed with the Hamilton Anxiety Rating Scale (HRSA), which consists of 14 items with a range of 0-56 (39). Higher scores on both scales reflects greater severity of symptoms.

### Bile Acid Processing

At the baseline visit, blood was collected via antecubital venipuncture regardless of fasting/fed status or time of day. After 20 minutes to allow for clotting, the blood was centrifuged at 4LC for 10 minutes and the serum was then immediately frozen at -80LC until ready for metabolomic analysis. BAs were quantified by ultra-performance liquid chromatography triple quadrupole mass spectrometry (Waters XEVO TQ-S, Milford, USA) by applying targeted metabolomics protocols and profiling protocols established in previous studies (40–42). Using a targeted approach, we originally quantified 36 primary and secondary, conjugated and unconjugated BAs. Our previous work reporting bile acid distributions in this dataset (31), demonstrated a significant association between the severity of depression and anxiety and the concentration of CDCA, along with its secondary BAs such as LCA, but not with CA or its secondary BAs. Therefore, for the current analysis incorporating neuroimaging, we focused on CDCA and LCA.

### Neuroimaging Image acquisition

MRI scans were performed up to one week prior to the baseline visit, as described previously (33). All imaging was conducted using a 3-T Siemens TIM Trio (Siemens Medical Systems, Erlangen, Germany). Resting-state fMRI was performed with eyes-open for 7.4 mins using the following protocols: a Z-SAGA sequence (43) to recover areas affected by susceptibility artifact; 150 volumes; 30 axial slices; voxel resolution= 3.4×3.4×4 mm^3^; matrix= 64×64; repetition time (TR)= 2950ms; echo time (TE) = 30ms. A high-resolution anatomical T1-weighted image was also acquired using a magnetization prepared rapid acquisition gradient-echo sequence (MPRAGE; TR= 2600ms, inversion time = 900ms, TE=3.02ms; flip angle = 8°; voxel resolution= 1.0×1.0×1.0 mm; number of slices = 176; matrix = 224×256).

### MRI data preprocessing

Image analysis used the AFNI (Analysis of Functional NeuroImages) software package; preprocessing was performed with the standard pipeline implemented in the AFNI package (44–46). All preprocessing steps and data quality assurance criteria are consistent with our prior work on this dataset (35,36). The functional image was aligned with the corresponding T1-weighted anatomical image, confirmed visually. The time series of rs-fMRI data were despiked and corrected for motion and slice-time acquisition. Scans with head motion exceeding 2 mm in any direction were excluded. The mean ± standard deviation of head motion, assessed as framewise displacement (FD) (47), were 0.1695 ±0.097. Furthermore, the time series were band-pass filtered (0.01<f<0.1Hz), and at the same time, the remaining effects of the noise signal, including residual head motion inferences, signals from the local white matter and CSF, were removed (48). The residual time series were then spatially smoothed up to 8 mm full-width at half-maximum (FWHM). Finally, the processed time series were normalized to standard Montreal Neurological Institute (MNI) space using the co-registration and anatomical image-based normalization parameters.

### Functional connectivity analysis

Due to our interest in the cognitive effects of bile acids, we focused on the ECN and DMN using a seed-based approach to assess their resting state functional connectivity (RSFC). Bilateral spheres of 5mm radius were used as seeds for the ECN (DLPFC seeds, x=±36, y=+27, z=29), and the DMN (PCC seeds, x=±7, y=-43, z=+33) based on the coordinates established by previous studies (49,50). The time series within these bilateral seeds were averaged, and Pearson’s correlation coefficient maps were generated for each individual patient. Subsequently, voxelwise correlation coefficients were then z-scored by Fisher’s r-to-z transformation. These z-values were used to determine the levels of functional connectivity for each seed.

### Statistical Analyses

The relationship between RSFC and bile acid concentrations was assessed through voxel-wise multiple linear regression analyses, with age and sex included as covariates. All results were presented using a family wise error (FWE) rate corrected α<0.05, equivalent to an uncorrected p<0.002, with a minimum cluster size of 492 mm^3^, which was demonstrated to be a reasonable approach in a previous study (46). Although use of a threshold more lenient than p<.001 has been asserted to result in unreliable findings in fMRI studies (51), we followed the guidance of Cox and colleagues, who have advocated for a balance between voxel and cluster thresholds for fMRI analyses (46,52), consistent with our prior analyses with this dataset (35,36). The determination of the cluster size for multiple comparison correction was estimated using simulated noise volumes, assuming a non-Gaussian spatial autocorrelation function (ACF) with 3dFWHMx, based on a Monte-Carlo simulation with 100,000 iterations, as implemented in 3dClustSim, with a cohort gray matter mask (53).

## RESULTS

Seventy-four patients had usable fMRI data and bile acid concentrations available for the analysis. Demographic and clinical characteristics of the sample are presented in **Table 1**.

**Table 1.** Clinical and Demographic Characteristics

| **Characteristic** | **M** | **SD** |
| --- | --- | --- |
| Age (yrs) | 38.5 | 11.0 |
| Baseline HAM-D | 19.1 | 3.0 |
| Baseline HAM-A | 15.8 | 3.9 |
|  | **<u>n</u>** | **<u>%</u>** |
| Sex |  |  |
| Male | 30 | 40.5 |
| Female | 44 | 59.5 |
| Race |  |  |
| Asian | 1 | 1.4 |
| Black | 12 | 16.2 |
| Multiple | 4 | 5.4 |
| Native American | 22 | 29.7 |
| White | 34 | 45.9 |
| Ethnicity |  |  |
| Hispanic | 26 | 35.1 |
| Non-Hispanic | 48 | 64.9 |
| Highest Education |  |  |
| < High school | 10 | 13.5 |
| High school or Trade school | 32 | 43.2 |
| University degree(s) | 32 | 43.2 |
| Employed Full-time |  |  |
| Yes | 34 | 45.9 |
| No | 40 | 54.1 |
| Lifetime major depressive episodes |  |  |
| 1 | 39 | 52.7 |
| 2 | 10 | 13.5 |
| ≥3 | 25 | 33.8 |
| Chronic episode (≥ 2 yrs)* |  |  |
| Yes | 17 | 23.0 |
| No | 57 | 77.0 |
| Anxiety disorder at baseline |  |  |
| Yes | 35 | 47.3 |
| No | 39 | 52.7 |

**Table 2.**
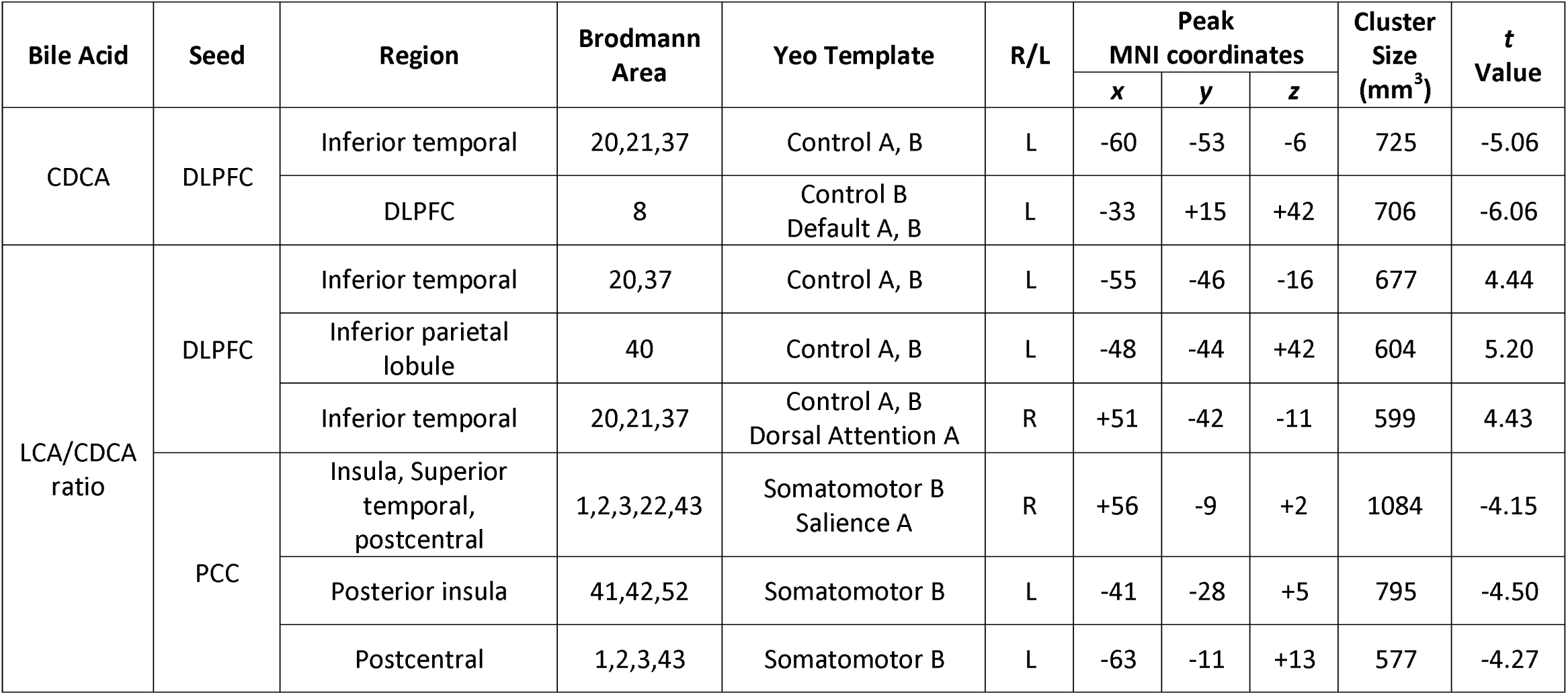
Resting state functional connectivity patterns significantly correlating with bile acid concentrations

## CDCA

CDCA levels demonstrated a strong significant inverse correlation (R^2^= .401, p<.001) with the RSFC between the DLPFC seeds and a left DLPFC region adjacent to the left-sided seed (**Fig. 1A**).

**Figure 1.**
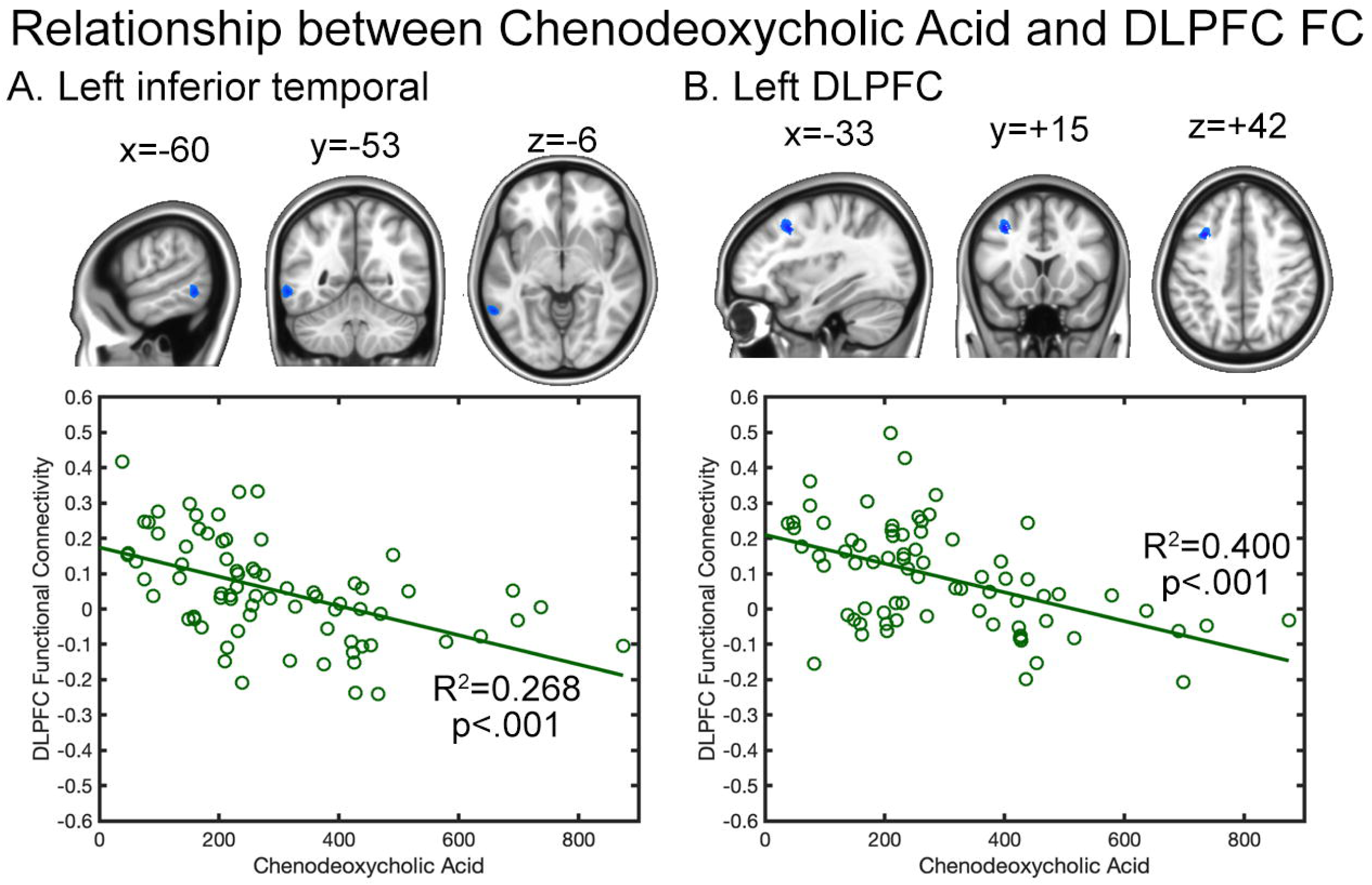
Strength of associations between brain regions with significant RSFC with the DLPFC seeds of the ECN and peripheral levels of bile acids. A. Chenodeoxycholic Acid (CDCA). CDCA levels are inversely correlated with the RSFC of the DLPFC seeds and another region in the left DLPFC that is part of the ECN. B. Lithocholic Acid (LCA). LCA levels are positively correlated with the with the RSFC of the DLPFC seeds and left inferior temporal cortex region of the ECN.

## LCA

LCA levels demonstrated a significant positive correlation (R^2^=.263, p<.001) with the RSFC between the DLPFC seeds and the left inferior temporal cortex (**Fig. 1B**).

## LCA/CDCA Ratio

The ratio of LCA/CDCA demonstrated significant correlations with two components of the ECN: the inferior temporal cortex and parietal lobe. **Fig. 2A and 2C** presented the significant RSFC between the DLPFC seeds and the left (R^2^=.298, p<.001, **Fig. 2A**) and right (R^2^=.270, p<.001, **Fig. 2C**) inferior temporal cortex. **Fig. 2B and 2D** presented the significant RSFC between the DLPFC seeds and the left inferior parietal lobule (R^2^=.248, p<.001, **Fig. 2B**) and the left superior parietal lobule (R^2^=.278, p<.001, **Fig. 2D**).

**Figure 2.**
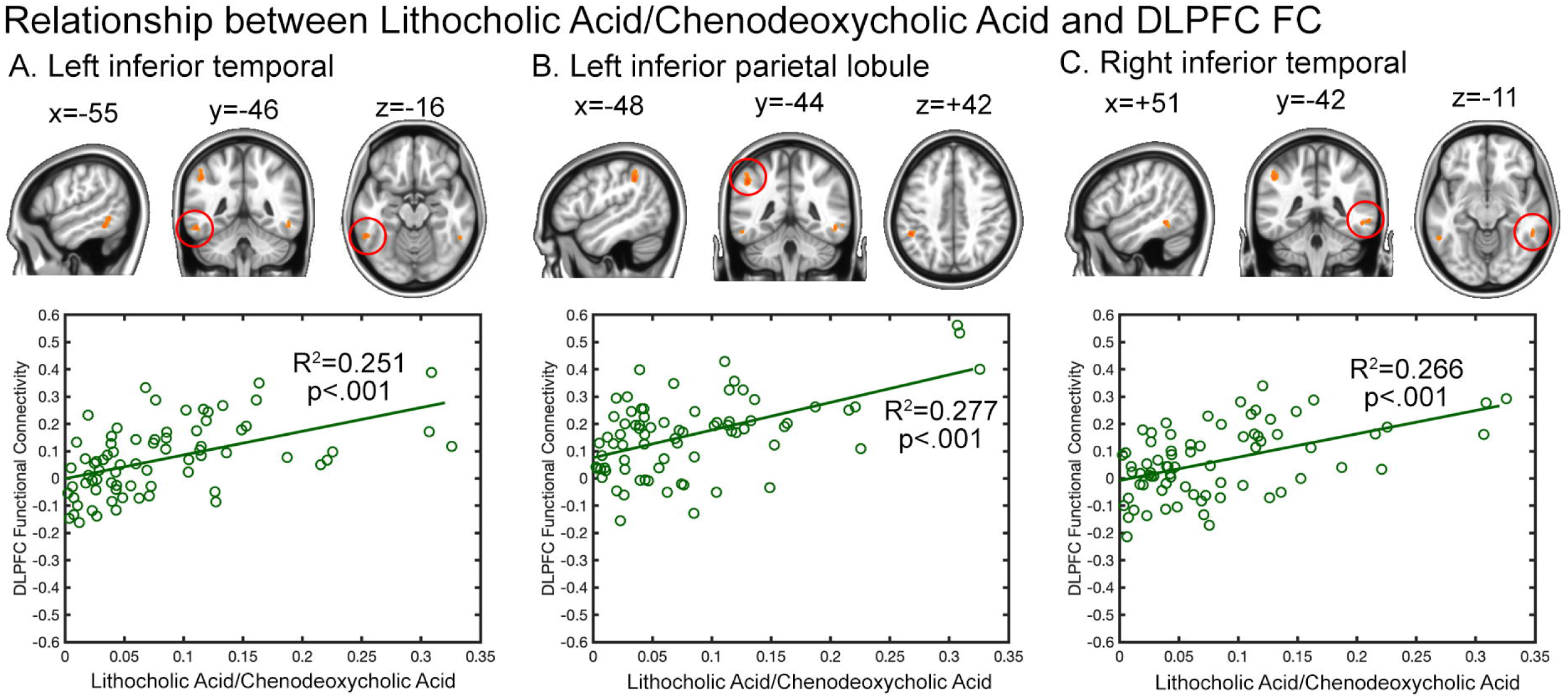
Strength of associations between brain regions with significant RSFC with the DLPFC seeds of the ECN and the ratio of LCA/CDCA. LCA/CDCA ratios are positively correlated with the RSFC of the DLPFC seeds with the left (A) and right (C) inferior temporal cortex regions of the ECN. LCA/CDCA ratios are positively correlated with the RSFC of the DLPFC seeds with the left superior parietal cortex of the ECN (B) and left inferior parietal cortex (D).

For PCC seeding for DMN functional connectivity, the ratio of LCA/CDCA demonstrated significant inverse correlations with several sections of the bilateral insula. **Fig. 3A and 3B** present the significant RSFC between the PCC seeds and the right (R^2^=0.270, p<.001, **Fig. 3A**) and left (R^2^=.275, p<.001, **Fig. 3B**) mid-insula regions. **Fig. 3C and 3D** presented the significant RSFC between the PCC seeds and the right (R^2^=0.259, p<.001, **Fig. 3C**) and left posterior insula (R^2^=0.256, p<.001, **Fig. 3D**).

**Figure 3.**
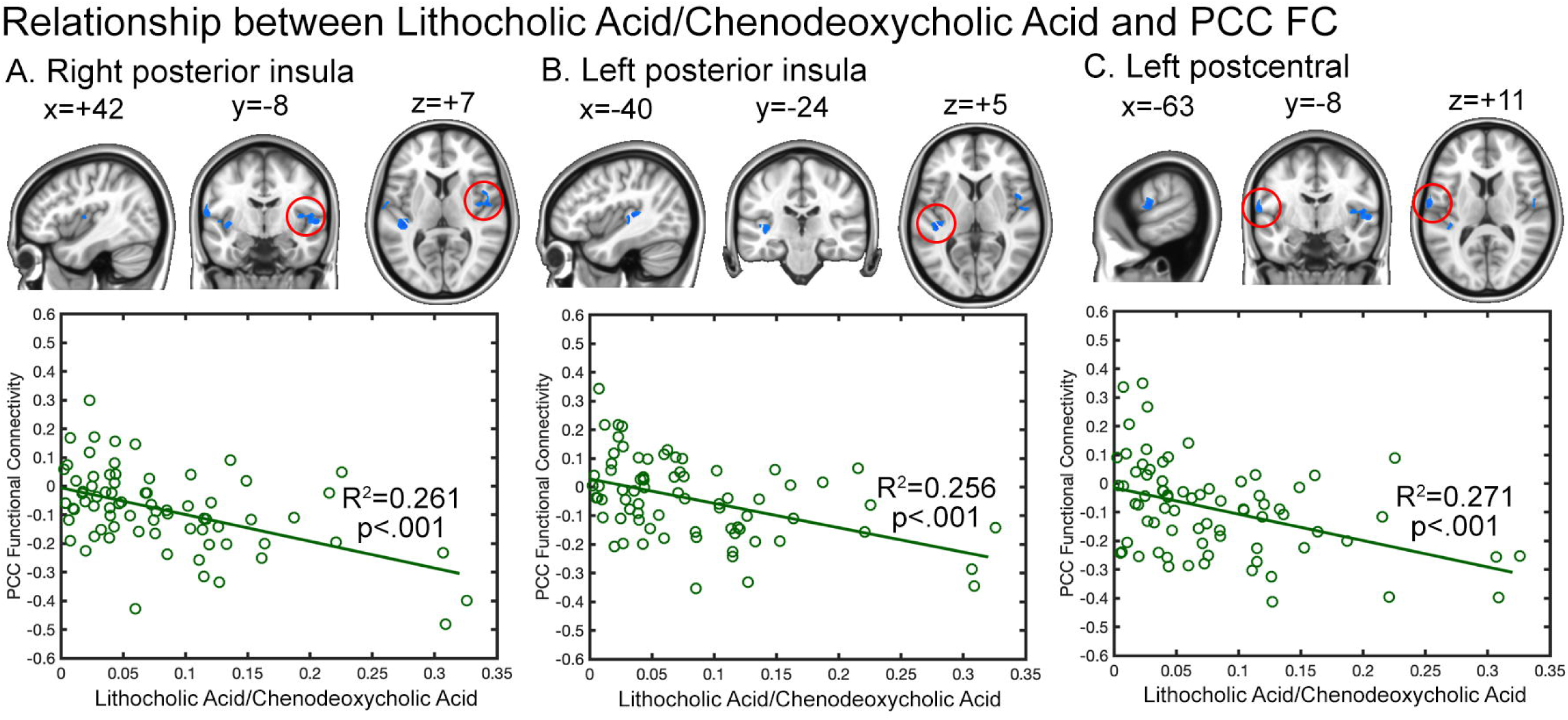
Strength of associations between brain regions with significant RSFC with the PCC seeds of the DMN and the ratio of LCA/CDCA. LCA/CDCA ratios are inversely correlated with the RSFC of the PCC seeds with the right (A) and left (B) mid-insula cortex, as well as the right (C) and left (D) posterior insula cortex.

### Clinical correlations

HRSD_17_ and HAMA scores did not correlate with either the CDCA level nor the CDCA-associated RSFC network. There was a weak association between the LCA/CDCA ratio with HAMA score (R^2^ =.054, p=.046) but no association with HAMD score. Neither the HAMA nor HRSD_17_ scores significantly correlated with the RSFC patterns associated with the LCA/CDCA ratio.

## DISCUSSION

The results of this study demonstrate significant associations between certain BA levels and activity within two established brain networks important for cognitive functioning among patients with MDD. The strongest association was found between the level of CDCA, a primary BA, and lower RSFC within the DLPFC. Conversely, levels of LCA, the microbiome-derived metabolite of CDCA, were positively correlated with the RSFC between the DLPFC seed and left inferior temporal cortex, which is a component of the ECN. The LCA/CDCA ratio more thoroughly revealed the relationship between these BAs and ECN network activity. Higher LCA/CDCA ratios were positively correlated with RSFC within core components of the ECN: DLPFC-bilateral inferior temporal cortex and DLPFC-left superior parietal cortex. These results are particularly striking because a whole-brain connectivity analysis was used; only the RSFC between regions within the ECN produced significant associations with BA levels. The specificity of these results strongly implicates the impact of BAs on cognitive systems in the brain.

There are intriguing relationships between the findings in this study of patients with MDD and the more numerous studies of the role of BAs in patients with AD. One notable study in AD patients with structural MRI data examined 15 BAs and found that the ratio of glycolithocholic acid (GLCA, the glycosylated form of LCA) to CDCA was associated with reduced cortical thickness in the same ECN regions that emerged in our analysis, specifically bilateral frontal, parietal, and temporal lobes (28). The GLCA/CDCA ratio was also associated with reduced hippocampal volume and reduced glucose metabolism in the temporal and parietal lobes, as assessed by (^18^F)FDG-positron emission tomography. Unfortunately, the LCA bile acid did not pass the quality control in pre-processing in that study and could not be analyzed (28).

AD pathophysiology implicates the accumulation early in the disease of extracellular amyloid-β (Aβ) containing plaques in the core areas of the DMN, specifically medial frontal cortex and medial parietal cortex (54). Combined with the observation that abnormal levels of Aβ_42_ in cerebrospinal fluid are associated with reduced connectivity between the DMN and ECN in older adults without brain atrophy, a hypothesis has been proposed that network failure in AD begins with the posterior DMN components (i.e., medial parietal cortex) even before amyloid plaques can be identified via imaging (55). Furthermore, longitudinal changes in Aβ pathology have been associated with fMRI-identified impairments in connectivity in the DMN and salience networks in individuals with preclinical AD, though these changes were not associated with changes in clinical cognitive performance (55). Notably, the current study found reduced connectivity between the DMN and insula regions was associated with the LCA/CDCA ratio.

Taken together, these results suggest the possibility that LCA/CDCA ratios in patients with MDD impact the same networks as Aβ plaques in patients with AD. Whether these overlapping findings represent two distinct processes acting on DMN functional connectivity, or whether these BAs are causally connected to Aβ plaque development should be a target of future research.

BAs have complex interactions with inflammatory processes and neurotransmitters within the CNS that may contribute to the neuroimaging results observed in this analysis. Although the blood-brain barrier (BBB) tightly regulates the passage of substances into the brain, evidence suggests that BAs can enter the brain through specific transport mechanisms or by passive diffusion when there is an alteration in BBB permeability (56). Inside the brain, BAs may interact with nuclear receptors such as the farnesoid X receptor (FXR) and G protein-coupled bile acid receptor 1 (GPBAR1, also known as TGR5), which are expressed in various brain regions (57). For example, some BAs, through their interaction with receptors like TGR5, can exert anti-inflammatory effects by inhibiting the production of pro-inflammatory cytokines and promoting the expression of anti-inflammatory cytokines in glial cells. This modulation of the inflammatory response could protect neurons from inflammatory damage, thereby preserving cognitive functions (58). BAs have been shown to influence the synthesis, release, and reuptake of neurotransmitters such as glutamate, GABA, and dopamine (59,60). These neurotransmitters are critical for cognitive processes including learning, memory, and decision-making.

The only clinical correlate with bile acid concentrations in this analysis was a weak positive association between the LCA/CDCA ratio and HAMA scores. An earlier analysis of the PReDICT sample, which was not limited to patients with usable fMRI data, found that among 208 patients at pre-treatment baseline, CDCA levels were significantly lower in patients with more severe depression symptoms and in those who were highly anxious (31). In addition, LCA levels and the LCA/CDCA ratio were significantly higher in the more anxious participants. Lower power from the reduced number of subjects (n=74) in the current analysis may have contributed to the minimal clinical associations identified here. However, despite the smaller sample, the significant association between LCA/CDCA and HAMA scores was replicated in the current analysis. In this light, it is notable that in the larger sample, an interaction analysis of the HRSD_17_ and HAMA scores indicated that differences in BA concentrations across patients correlated more strongly with anxiety symptoms than depressive symptoms (31).

This analysis has several strengths. The study utilized one of the few extant datasets that permits the correlation of peripheral metabolomic signals with brain activity in patients with psychiatric disease. The sample was medically healthy and all were treatment-naïve with respect to antidepressants, precluding a confounding role of antidepressants or anxiolytics on the neuroimaging data. The patient population was reasonably diverse in terms of gender, race, and ethnic composition. The major limitation of this study was the absence of a clinical measure of cognitive functioning, which was not included in the PReDICT study design. Consequently, we were unable to identify a clinical correlate that was associated with the bile acid levels or their impact on ECN connectivity. An additional limitation is the 7.4 minute duration of the resting state scan, which could have reduced the power and reliability of the functional connectivity analyses (61).

## CONCLUSIONS

In summary, the impact of LCA and CDCA levels on the RSFC of the ECN and DMN in MDD patients overlaps remarkably with the neuroimaging findings of the impact of BAs and Aβ plaques in patients with AD. It is possible that the previously identified increased risk for dementia diagnoses in patients with MDD in mid-life may be driven in part by the subset of MDD individuals with elevated LCA/CDCA ratios. Testing this hypothesis should be a target of future research. If verified, this model would point to the need to develop novel treatment interventions that target the microbiome to diminish the production of neurotoxic BAs as a potential mechanism for preventing or at least forestalling the development of AD.

The results of this study underscore the importance of examining the role of BAs, particularly those derived from gut microbiome metabolism, as a potential contributor to cognitive symptoms in patients with MDD, and their potential role for presaging the development of cognitive disorders later in life.

